# Interactive mapping of language and memory with the GE2REC protocol

**DOI:** 10.1101/2019.12.24.888040

**Authors:** Sonja Banjac, Elise Roger, Emilie Cousin, Marcela Perrone-Bertolotti, Célise Haldin, Cédric Pichat, Laurent Lamalle, Lorella Minotti, Philippe Kahane, Monica Baciu

**Author notes:** **Corresponding Author:** Monica Baciu, LPNC, UMR CNRS 5105, BSHM, BP 47, 38040 Grenoble Cedex 09, France.

## Abstract

Previous studies have highlighted the importance of considering cognitive functions in a dynamic and interactive perspective and multiple evidence was brought for a language and memory interaction. In this study performed in healthy participants, we developed a new protocol entitled GE2REC that interactively accesses the neural representation of language-and-memory network. This protocol consists of three runs related to each other, providing a link between tasks, in order to assure an interactive measure of linguistic and episodic memory processes. GE2REC consists of a sentence generation (GE) in auditory modality and two recollecting (2REC) memory tasks, one recognition performed in visual modality, and another one recall performed in auditory modality. Its efficiency was evaluated in 20 healthy volunteers using a 3T MR imager. Our results corroborate the ability of GE2REC to robustly activate a fronto-temporo-parietal language network as well as temporal mesial, prefrontal and parietal cortices during encoding and recognition. GE2REC is useful because: (a) requires simultaneous and interactive language-and-memory processes and jointly maps their neural basis; (b) explores encoding and retrieval, managing to elicit activation of mesial temporal structures; (c) is short and easy to perform, hence being suitable for more restrictive settings, and (d) has an ecological dimension of tasks and stimuli. Overall, GE2REC can provide valuable information in terms of the practical foundation of exploration language and memory interconnection.

## 1. Introduction

As suggested by recent studies, the base of proper cognitive functioning is the dynamic interaction between different neuropsychological domains (Kellermann et al., 2016). Language and memory, two fundamental cognitive functions, are not isolated abilities but instead are embedded in each other and in other cognitive processes, contrary to views considering them as independent modules (Newell, 1990). There is cognitive evidence suggesting that memory and language influence each other more than previously thought (Huettig and Janse, 2016; Moscovitch, Cabeza, Winocur, & Nadel, 2016; Vogelzang et al., 2017). The relation between language and working memory has already been largely supported (Vogelzang et al., 2017; Määttä et al., 2014; Kowialieswski & Majerus, 2019). At the same time, the relation between language and other memory domains is still under investigation. For example, the language and memory tie is visible in autobiographic memory. Language influence is evident in the studies showing that the access to autobiographical memories is better if subjects are using the same language during memory formation and remembering (Larsen, Schrauf, Fromholt, & Rubin, 2002; Marian & Neisser, 2000). On the other hand, memory functioning can manifest in language production through the usage of different linguistic constructions (Park, St-Laurent, McAndrews, & Moscovitch, 2011). Several cognitive frameworks posit the commonalities between language and memory function in terms of their underlying brain systems (Ullman, 2001; Ullman, 2004; Pinker and Ullman, 2002). Nevertheless, how this language and memory interaction is supported by common neural systems remains largely unexplored.

The aim of the present study situated within this framework is to continue this line of reasoning by exploring the interaction of language and episodic memory, or as we will refer to it henceforth language-and-memory and proposing a protocol that would allow to map out the neural representations of this network. Several categories of arguments underline the interaction between language and memory networks and processes. First, there is anatomical evidence that provides the neural basis of language-and-memory network. Namely, the central regions of episodic memory represented by lateral and medial temporal structures, notably hippocampus and entorhinal cortex, have connections with the structures that are engaged in language. Specifically, there is a direct inter-hippocampal pathway that projects inferior temporal cortex (BA 37) connected with the inferior visual system to hippocampus and transmits from entorhinal to temporal association cortex, temporal pole and prefrontal cortex. Also, there is a polysynaptic pathway that projects posterior parietal association cortex (BA 7) and the temporal cortices (BA 40, 39, 22) over parahippocampal gyrus to the entorhinal area, and conducts the hippocampal outputs via anterior thalamic nucleus to posterior and anterior cingulate and retrosplenial cortices (BA 29 and 30) (Duvernoy, 2005; Tracy & Boswell, 2008). In accordance with these findings, studies have also shown that certain fronto-temporal fiber tracks are engaged in both language and memory functions. Namely, the uncinate fascicle (UF), connecting the anterior temporal and orbito-frontal area, has primarily been found to supports episodic encoding and retrieval (Diehl et al., 2008), but has also been recognized as a part of the naming network (Duffau, 2014; Kucukboyaci et al.,2014; McDonald et al., 2008). While the inferior fronto-occipital fascicle (IFOF) that was known as a part of ventral semantic stream (Duffau, 2014), has actually been discovered to be associated with amodal semantic processing, noetic and subsequently auto-noetic consciousness (Moritz-Gasser, Herbet, & Duffau, 2013) and verbal memory performances (McDonald et al., 2008). Moreover, in addition to classical perisylvian network, functional connectivity research on associative-semantic network has also identified the second subsystem encompassing ventral occipito-temporal transition zone, intraparietal sulcus and hippocampus which was assumed to serve as a mediator between general inner representations and other cognitive systems such as the episodic memory (Vandenberghe et al., 2013).

Furthermore, there are studies that acknowledged the interactive relationship of language and memory functions and explored different aspects and mechanisms of their bond. Namely, it has been proposed that language usage is highly dependent on declarative memory system, and that this interaction is being accomplished through hippocampus (Duff & Brown-Schmidt, 2012). Moreover, it has been shown that the language-and-memory dynamics change over the course of life. For example, it has been proposed that there is a common underlying mechanism in the development of language dominance and material specificity of mesio-temporal structures (Weber, Fliessbach, Lange, Kügler, & Elger, 2007) as well as that hippocampal structure may contribute to early language ability beyond its contribution to general intellectual ability (Lee et al., 2015), and it was found that language skills are linked to children’s recall abilities (Klemfuss, 2015). While it was shown in elderly, that the lexical production is based on higher usage of semantic memory which was observed through the changes in the underlying network connectivity between inferior frontal and medial temporal cortex (Hoyau et al., 2018).

Additional arguments on language and memory interaction come from studies on disorders such as post-stroke aphasia (Schuchard & Thompson, 2014) or conditions that cause auditory hallucinations (Ćurčić-Blake et al., 2017) in which language regions are impaired, but the symptoms also manifest in the memory domain. Nevertheless, this dynamical relation between language and memory is most apparent and hence studied in temporal lobe epilepsy (TLE) that represents 70-80% of epilepsy in adults (Jaimes-Bautista, Rodríguez-Camacho, Martínez-Juárez, & Rodríguez-Agudelo, 2015) and is characterized by seizures induced by a regional dysfunction, the epileptic zone (EZ), located in temporal regions. As language and memory networks converge towards integrative hubs that mainly stem from the left temporal lobe (Battaglia, Benchenane, Sirota, Pennartz, & Wiener, 2011; Mesulam, 2000), joint language-and-memory deficits are common in these patients, even greater if the EZ is located mesially (Perrone-Bertolotti, Zoubrinetzky, Yvert, Le Bas, & Baciu, 2012; Zalonis et al., 2017). In 30-60% of left TLE (LTLE) patients, the verbal memory declines (Bell, Lin, Seidenberg, & Hermann, 2011) as well as long-term memory (Tramoni-Negre, Lambert, Bartolomei, & Felician, 2017). At the same time, studies suggest that between 40 and 55% of patients with LTLE and 36% of those with right TLE (RTLE) show naming deficits (Bartha-Doering & Trinka, 2014; Zhao et al., 2014). Additionally, TLE patients with hippocampal sclerosis (HS) have been found to have worse naming performance than those without it (Davies et al., 1998) and the volume of left hippocampus has been found to significantly predict verbal fluency and naming (Alessio et al., 2006). Generally, studies with TLE patients show that hippocampus has a vital role in retrieving lexically and semantically associated words (Bonelli et al., 2011; Hamamé, Alario, Llorens, Liégeois-Chauvel, & Trébuchon-Da Fonseca, 2014) and that it is active during language comprehension tasks, as well as its neighboring structures (Meyer et al., 2005). Importantly, Piai et al. (2016) have found that the same hippocampal neuronal computations estimated using theta waves can also be identified during language tasks, which is why it was proposed to incorporate hippocampus into language network (Covington & Duff, 2016). These findings underscore again the strong interactions between language and memory.

Moreover, nearly 35% of TLE patients are refractory to drugs (Harroud, Bouthillier, Weil, & Nguyen, 2012) and curative surgery to remove or inactivate the EZ remains the only solution to stop seizures (Schoenberg, Werz, & Drane, 2011; Téllez-Zenteno, Dhar, & Wiebe, 2005). As temporal regions are crucial for language and memory, surgery in these regions must be preceded by detailed preoperative mapping of these functions, to avoid possible postoperative deficits (Baxendale, Thompson, Harkness, & Duncan, 2006; Drane & Pedersen, 2019; Gleissner, Helmstaedter, Schramm, & Elger, 2004; Hamberger, 2007, 2015; Helmstaedter, Kurthen, Lux, Reuber, & Elger, 2003; for a review see Sherman et al., 2011). Importantly, seeing that language and memory mechanisms and substrates are highly interconnected (Huettig & Janse, 2016; Vogelzang et al., 2017; Tracy & Boswell, 2008), their assessment should be performed in interaction and interplay.

Despite the substantial evidence supporting the idea of close interconnection of language-and-memory, one can question the advantages of exploring them interactively. First of all, present language models (e.g. Duffau, Moritz-Gasser & Mandonnet, 2014; Hickok & Poeppel, 2007; Indefrey & Levelt, 2004; Price, 2012; except Ullman, 2004) do not incorporate mesial temporal structures that have been shown to contribute to language processing (Alessio et al., 2006; Bonelli et al., 2011; Hamamé, et al., 2014) overlooking thus to acknowledge the necessity for a dynamic interaction between the domains in order to achieve successful cognitive functioning (Kellermann et al., 2016). Developing a neurocognitive model that encompasses both language and memory would help understanding more profoundly their interaction, but also to explore and understand brain plasticity better. Namely, in TLE patients there is a possibility of exploring cerebral reorganization of language and memory networks that have double origin, chronic epilepsy and surgery and can be described at inter and intra-hemispheric level (Baciu & Perrone-Bertolotti, 2015; Berl et al., 2014; Dupont et al., 2000; Powell, Parker, et al., 2007; Powell, Richardson, et al., 2007; Rosenberger et al., 2009; Sidhu et al., 2013; Springer et al., 1999). Inter-hemispheric reorganization is manifested in the recruitment of non-dominant hemisphere regions, whereas intra-hemispheric reorganization consists of recruitment of regions in the dominant hemisphere, initially non-specialized for language or memory (Cousin, Baciu, Pichat, Kahane, & Le Bas, 2008). Overall, inter and intra-hemispheric reorganization of cognitive networks is based on modulations of reciprocal inter-hemispheric excitatory and inhibitory interactions that maintain the hemispheric specialization in normal condition (Tzourio-Mazoyer et al., 2017). Importantly, the reorganization of language network can depend on regions that have not classically been considered a part of that network (Tracy & Boswell, 2008) such as hippocampus (Baciu & Perrone-Bertolotti, 2015). Another practical importance of exploring language-and-memory would be the already mentioned pre-surgical evaluation, since the reorganization is not necessarily effective (Golby et al., 2002; Powell et al., 2007; Sidhu et al., 2013). There is a number of previous studies that have shown that visualizing the reorganization of language (Rosazza et al., 2013) or memory (Bonelli et al., 2010; Massot-Tarrús, White, & Mirsattari, 2019) networks before the surgery can predict the postoperative decline of these functions. However, more importantly, language fMRI activation has been found to predict verbal memory postoperative outcome (Binder et al., 2008; Binder et al., 2010; Labudda, Mertens, Aengenendt, Ebner, & Woermann, 2010). Considering it together with the evidence of high interconnection between language and memory mechanisms and substrates (Tracy & Boswell, 2008), it is reasonable to conclude that their assessment should be performed in interaction and interplay. Functional MRI is a method of choice to assess cerebral reorganization (Abbott, Waites, Lillywhite, & Jackson, 2010; Detre, 2006; Sabsevitz et al., 2003). Even though fMRI is currently regarded as an efficient tool for preoperative assessments of cortical regions for purposes of resection optimization (Beers & Federico, 2012; Binder, 2011), there is no consensus for the most appropriate protocol and paradigm to determine language and memory brain lateralization and localization (Benjamin et al., 2018; Perrone-Bertolotti et al., 2015). Certain authors have proposed fMRI protocols that encompassed both language and memory tasks (Aldenkamp et al., 2003; Deblaere et al., 2002). However, they examined these two functions separately, concluding afterwards about their interconnection. There is also an important issue of protocols’ ecological validity (Mayer & Murray, 2003) that is often neglected. Bearing this goal in mind, one important characteristics of fMRI protocols should be their proximity to everyday experiences. In addition to that, seeing that identified reorganization is not necessarily effective (Golby et al., 2002; Powell et al., 2007; Sidhu et al., 2013), we particularly underscore the importance of correlating brain activity with behavioral and cognitive performances, to evaluate reorganization efficiency. An efficient pattern of reorganization indicates that brain restructuring is associated with normal cognitive performance. Brain activation observed during language and memory mapping is largely determined by the nature of the task used to explore them (Baciu & Perrone-Bertolotti, 2015; Bradshaw, Thompson, Wilson, Bishop, & Woodhead, 2017; Perrone-Bertolotti et al., 2015). Generally, language tasks should map a network encompassing inferior frontal region (pars triangularis, opercularis and orbitalis), insula, superior, medial and inferior temporal gyri, supramarginal guyrs, angular gyrus, supplementary speech area and occipito-temporal area (Benjamin et al., 2017; Labache et al., 2018; Price, 2012; Vigneau et al., 2006) with Crus 1 and 2 and IV, V, VI, VIIAt and VI lobules of cerebellum (Keren-Happuch, Chen, Ho, & Desmond, 2014; Price, 2012; Stoodley & Schmahmann, 2018). In addition, hippocampus, entorhinal, perirhinal and parahippocampal cortices together with amygdala, cingulum, lateral orbito-frontal gyrus, medial prefrontal cortex, superior and inferior parietal area are specifically involved in encoding and/or retrieval process during long-term memory evaluation (Battaglia et al., 2011; de Vanssay-Maigne et al., 2011; Diana, Yonelinas, & Ranganath, 2007; Ranganath & Ritchey, 2012; Spaniol et al., 2009).

In the present study, we present and evaluate an original fMRI protocol entitled GE2REC with the intention to map language-and-memory network in a concise and robust fashion. GE2REC consists of a sentence generation with implicit encoding (we abbreviate it GE) in auditory modality and two recollection (2REC) memory tasks, a recognition (we abbreviate it RECO) performed in visual modality, and a recall of sentences (we abbreviate it RA), performed in auditory modality. The GE and RA runs are designed to activate intermixed language-and-memory network by engaging episodic memory encoding and retrieval respectively. It is important to note that there is a change of modality between GE (audio) and RECO tasks (visual) in order to enhance the access to episodic memory and correspondingly the activation of hippocampal structures. The originality of the current protocol is reflected in the fact that it (a) explores the language-and-memory in interaction, (b) explores both encoding and retrieval (via both recognition and recall), (c) enhances hippocampal recruitment by engaging episodic memory, d) is easy to perform and has a short duration, and (e) has a natural, ecological dimension, particularly the RA task based on everyday life experiences.

## 2. Material and Methods

### 2.1. Participants

Twenty-one right-handed healthy volunteers aged between 18 and 29 years (M=21, SD= 3.3; 9 females), without neurological and psychiatric deficits were included in this study. All participants were French native speakers and had normal or corrected-to-normal vision. One participant was excluded from the fMRI analyses due to the high amount of artifacts in the data. This clinical experimentation is governed by the French law (Jardé, Décret n°2016-1537 16/11/2016 from 17/11/ 2016). Ethical approval was granted by national and local competent authorities. The Ethic committee for the protection of persons has approved the project (CPP 09-CHUG-14; MS -14-102). All participants provided written informed consent to participate to study and they received the financial compensation for their participation.

### 2.2. Functional MRI (fMRI) assessment of language and memory

The experimental protocol was developed using E-prime software (Psychology Software Tools, Pittsburgh, PA). Before entering into the magnet, participants were explained the outline of the procedure. Importantly, they only received full description of the task in the GE run. For the 2REC runs they were only informed about the general outline of the tasks and how they should respond, while they remained uninformed about the actual content of the tasks. A schematic illustration of all tasks is presented in Figure 1.

**Figure 1.**
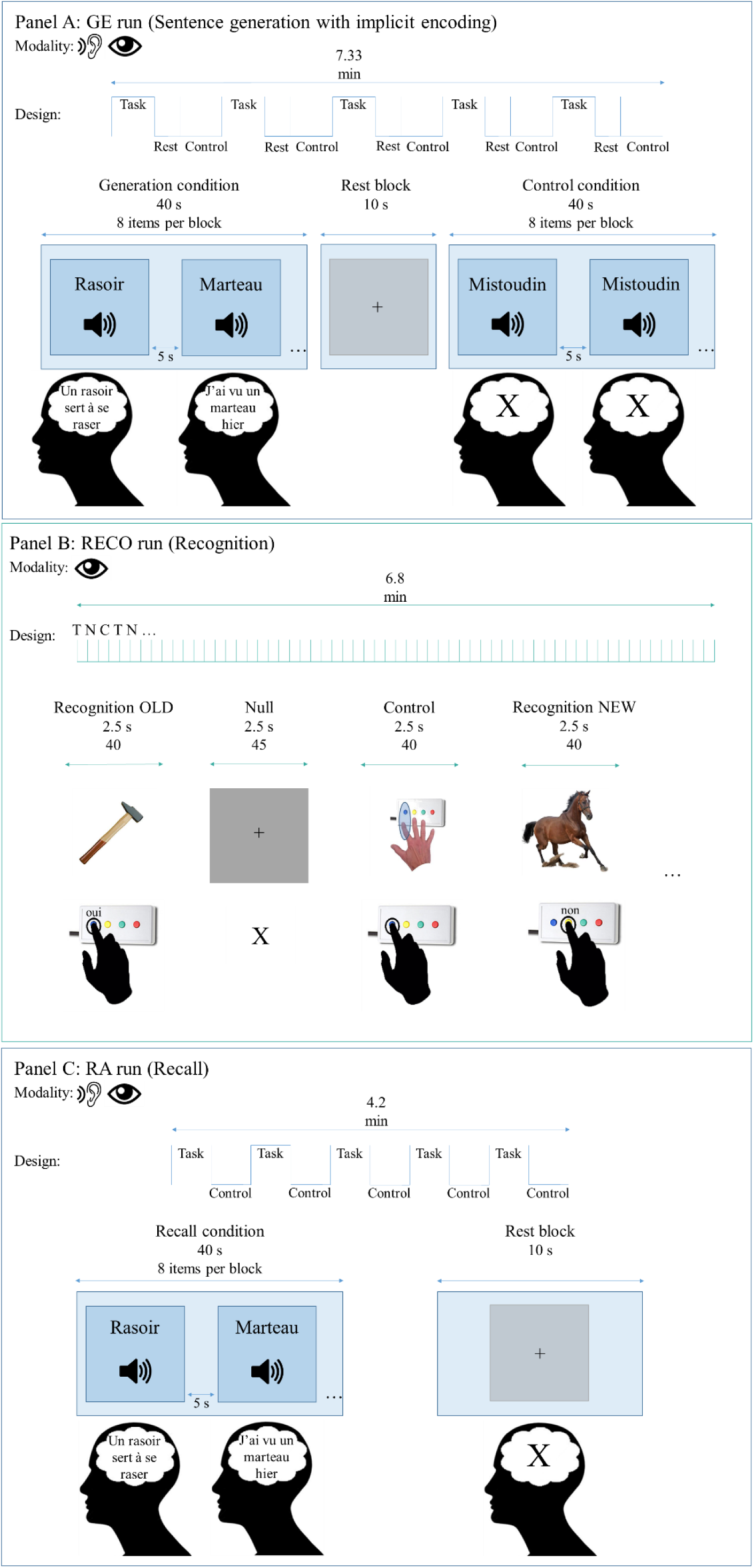
Schematic illustration of the GE2REC protocol. **Panel A:** GE (Sentence generation with implicit encoding) run with block-design. Items were presented in auditory modality during Task (word to generate sentences) and Control (pseudo-word) and in visual modality during Rest (central cross to fixate). Participants were required to covertly generate sentences during Task and to do nothing during Control. They fixated the cross during Rest. Examples of French items are shown *(rasoir=razor; marteau=hammer; mistoudin is a pseudo-word)*. **Panel B:** RECO (recognition) run with the event-related design. Items were presented in visual modality during Task (images to recognize), Control (images to be repeated) and Null events (central cross to fixate). Participants were required to recognize whether or not they have heard the object presented in the image and to reply by using the response box. During the Control, they were asked to press the button shown in the picture and to fixate the cross during the Null event. **Panel C:** RA (Recall) run with block-design. Items were presented in auditory modality during Task (word to recall sentences) and in visual modality during Rest (central cross to fixate). Participants were required to recall the sentences they generated in the GE run and to covertly repeat them. They fixated the cross during Rest.

#### 2.2.1. GE stimuli and task

During the GE run, the participants heard words through a headset and their task was to covertly generate a sentence for each word they heard. A block design was used, including task, control and rest conditions. Words corresponded to images from the French standardized naming test D080 (Metz-Lutz et al., 1991). The run included 5 task conditions of sentence generation performed in auditory modality (8 stimuli/condition, 40 words in total) and the inter-stimulus intervals (ISI) that lasted 5 s that were intended to provide enough time to generate a correct sentence. The participants were instructed to generate a sentence, after hearing a word, by using the word they heard and to continue repeating it until hearing the next word. The generation was performed covertly. The run also included 5 control periods (non-generation) in order to control for auditory activations during which we played a pseudoword 8 successive times, with 5 s ISI. The participants were required to just listen the pseudoword and to try not to talk covertly. The run also included 5 rest blocks in visual modality that were represented by a fixation cross displayed during 10 s, placed directly after the generation blocks in order to provide time for the hemodynamic response to come down. Participants were required to simply fixate the cross. The order of conditions was Task (Generation), Rest and Control. The total duration of the run was 7.3 min.

#### 2.2.2. RECO stimuli and task

During the RECO run performed in the visual modality, the participants were shown pictures on the screen and their task was to respond whether they heard the names of the objects in the images during the GE run. The event-related design was used, including pictures of the words participants heard in the previous task, pictures of the new objects, control images and rest condition. All presented images were real-life equals of the images from the DO80 database (Metz-Lutz et al., 1991)^1^. The run included 40 pictures of the words presented in the GE run (henceforth OLD). The participants were instructed to press the “yes” button on their response box that was in their dominant hand when they saw the image that corresponded to one of the words they heard in the previous run. Additionally, the run included 40 pictures of the words that were not presented in the GE run (henceforth NEW) and were matched with the OLD by the length and frequency. The participants were required to press “no” button on their response box when they saw the image that was showing the object whose name they did not hear in the previous run. The run also included 40 control images showing the button that needed to be pressed in order to control for the motor activations during button pressing. Furthermore, the run contained 45 null events represented by a fixation cross. All events were displayed during 2.5 sec and conditions were presented in pseudo-randomized order. The total duration of the run was 6.8 min. We employed the event-related rather than block design since the former has been shown to identify well the effects of successful encoding (Haag & Bonelli, 2013) and in order to avoid the prediction of stimuli.

#### 2.2.3. RA stimuli and task

During the RA run, the participants heard again through a headset the words they have heard in the GE run. Their task was to recall and covertly repeat the sentence they have generated for each word in the GE run. A block design was used, including task and rest conditions. The run included 5 task conditions of recall performed in the auditory modality (8 stimuli/condition, 40 words in total) with 5 s ISI. The participants were instructed to remember and repeat the sentence they have generated in the first GE run and to continue repeating it until hearing the next word. The recall task was done covertly. The run also included 5 rest blocks in visual modality that were represented by a fixation cross displayed during 10 s and participants were asked to simply fixate the cross. The total duration of the run was 4.17 min.

Since fMRI is highly sensitive to motion (Powell & Duncan, 2005), we have decided to use covert production in GE and RA runs. This is a commonly used version of production task (Black et al., 2017) that has been proven to provide reliable activation of language regions and lateralization (Benjamin et al., 2017; Haag & Bonelli, 2013).

### 2.3. MR Acquisition

Functional MRI was performed at 3T (Achieva 3.0T TX Philips Medical systems, NL) at IRMaGe MRI facility (Grenoble, France). The manufacturer-provided gradient-echo/T2* weighted EPI method was used for the functional scans. Forty-two adjacent axial slices parallel to the bicommissural plane were acquired in sequential mode (3mm thickness, TR = 2.5 s, TE = 30ms, flip angle = 82°, in-plane voxel size = 3 × 3 mm; field of view = 240 × 240 × 126 mm; data matrix = 80 × 80 pixels; reconstruction matrix = 80 × 80 pixels). Additionally, for each participant a T1-weighted high-resolution three-dimensional anatomical volume was acquired, by using a 3D T1TFE (field of view = 256 × 256 × 160 mm; resolution: 1 × 1 × 1 mm; acquisition matrix: 256 × 256 pixels; reconstruction matrix: 256 × 256 pixels).

### 2.4. Data processing

#### 2.4.1. Behavioral analyses

Based on the responses during the RECO run, we calculated behavioral performances during recognition task (%CR_RECO). As mentioned above, the encoding performance during GE was indirectly determined via recognition (RECO). On the basis of the %CR_RECO for old items, we identified those that were successfully encoded among all items presented during GE. Statistical analyses were performed using RStudio software version 1.1.456 (RStudio Team, 2016). All one-sample and paired *t* tests were computed with “t.test” function in the “stats” R package version 3.5.1 (R Core Team, 2018).

#### 2.4.2. Functional MRI analyses

The Analyses were performed using SPM12 (Welcome Department of Imaging Neuroscience, London, UK) running under Matlab R2015b (Mathworks Inc., Sherborn, MA, USA).

##### • Pre-processing steps

Functional MRI data was first pre-processed along several steps. First, functional volumes were time-corrected with the mean image as the reference slice in order to correct artifacts caused by the delay of time acquisition between slices. Thereafter, all time-corrected volumes were realigned to correct the head motion. The T1-weighted anatomical volume was co-registered to mean images obtained through the realignment procedure, normalized to MNI (Montreal Neurological Institute) space and smoothed by an 8 mm FWHM (Full Width at Half Maximum) Gaussian kernel. Noise and signal drift were removed by using a high-pass filter (1/128 Hz cutoff). In the next step, these spatially pre-processed volume, were included into statistical analyses according to experimental conditions.

##### • Functional MRI statistical analyses

We evaluated GE and RA runs by analyzing them as a block design, while encoding (we abbreviate it ENCO) was analyzed using the GE run but as an event-related design by selecting only those GE items that were correctly recognized during the RECO run. In the same vein, the recognition was evaluated by analyzing the RECO run as event-related, using only the correctly recognized items and comparing it with RECO control condition, as well as comparing correct recognition of OLD and NEW items. Statistical parametric maps were generated from linear contrasts between the HRF parameter estimates for the different experimental conditions. The whole brain effects of interest were firstly evaluated at an individual level (first-level): (1) effect of language by comparing sentence generation and control; (2) effect of memory encoding by comparing the correctly encoded items during GE with the baseline; (3) effects of memory recognition by comparing correctly recognized old items with the control; (4) effects of memory recognition by comparing correctly rejected new items with the control; (5) differences in recognition by comparing recognition of old and new items and (6) effects of memory recall by comparing sentence repetition with the baseline. Six movement parameters obtained by realignment corrections were included as noise (regressors of non-interest).

For the second-level group analysis, individual contrasts were entered into a one-sample *t* test and activations were reported at a *p* < .05 significance level with the FWE corrected (T_GE_ > 6.5; T_RECO_ > 6.82; T_RA_ > 6.54) and threshold of 5 voxels (k > 5) for all effects except the memory encoding effect. Bearing in mind that a parametric modulation was performed and that fMRI acquisition of medial temporal lobe (that we were specifically interested) can be affected by geometric distortions and signal loss (Haag & Bonelli, 2013; Powell & Duncan, 2005), for the memory encoding effect we employed a p < .001 uncorrected significance level and threshold of 5 voxels (k > 5) with T_ENCO_ > 3.58.

## 3. Results

### 3.1. Behavioral results

Inspection of the performance during the RECO run showed that participants correctly recognized on average 72.62% (SD = 10.2) of old items and correctly rejected on average 87.87% (SD = 7.36) of new items. The correct recognition of old items and the correct rejection of new items were both above the chance level (*t*(20)_OLD_ = 10.16, p < .001; t(19)_NEW_ = 23.02, *p* < .001). Moreover, paired t test demonstrated that the recognition of old items (M_RT_OLD_ = 0.97; SD = 0.07) was faster (*t*(19) = −5.51, *p* < .001) than the rejection of the new ones (M_RT_OLD_ = 1.1; SD = 0.07).

### 3.2. Functional MRI

#### 3.2.1. Sentence Generation with implicit encoding (GE)

Comparison task vs. control showed a large cerebral substrate recruited by the GE task. Results are presented in Panel A of Figure 2, and activation peaks are mentioned in Table 1. Overall, the results reveal bilateral but left predominant activation of a large fronto-temporo-parietal network including left prefrontal, inferior frontal, and bilateral insula. In addition, activation of left superior temporal and bilateral middle temporal and superior temporal pole cortices were observed together with right precuneus and right cerebellum Crus 1 and VI.

**Figure 2.**
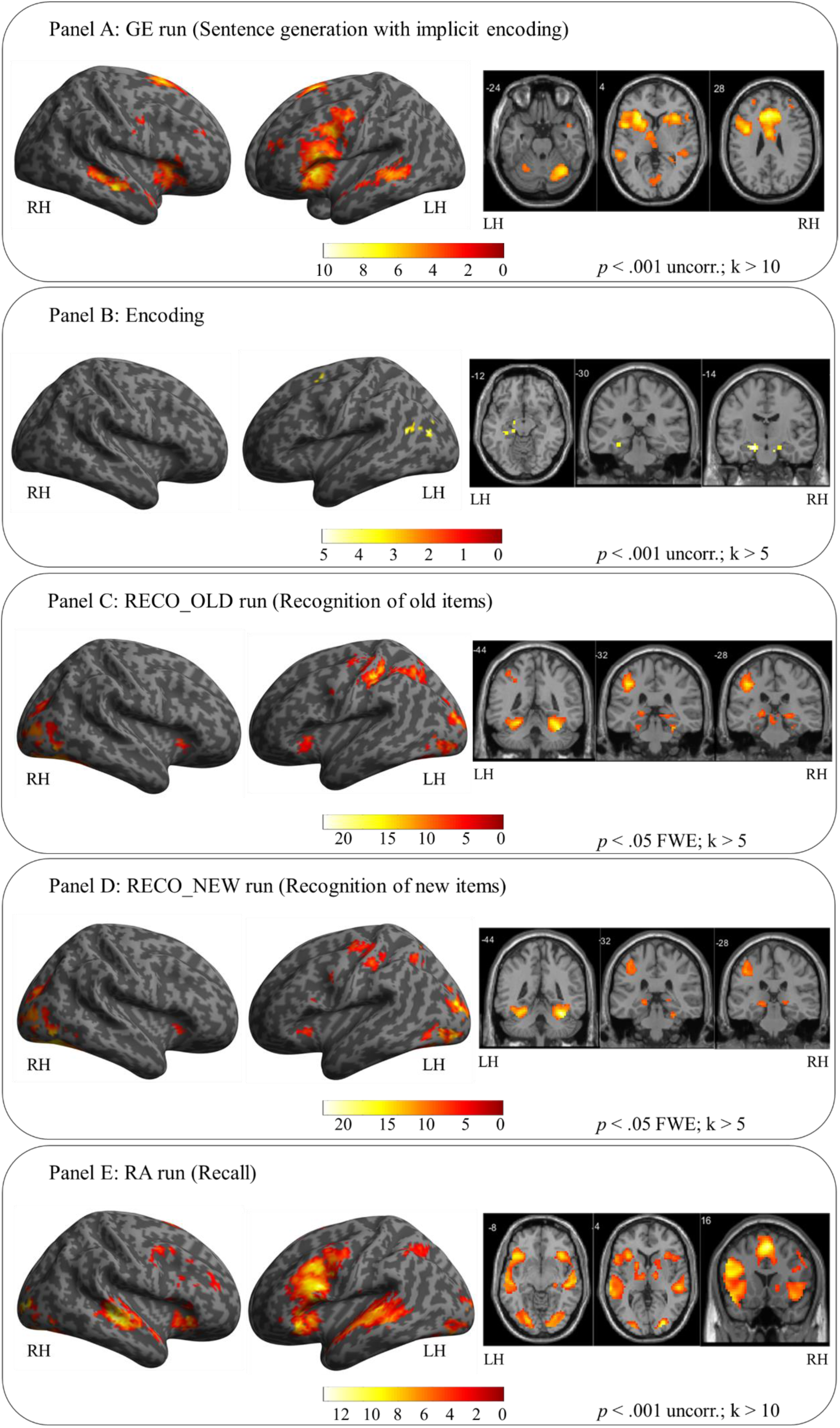
Illustrative overview representation of global activation obtained for sentence generation (panel A), encoding (panel B), recognition of old (panel C) and new items (panel D) and the recall (panel E). Activations for each task were obtained at a group level (N=20 participants for all tasks except recognition of new items where N=19 were included due to a lack of responses of one participant). Activations were projected onto the lateral left and right views of surface rendering and 2D coronal and axial slices. The left (LH) and right (RH) hemispheres are indicated. The color scale indicates the T value of the activation. The GE and RA runs, as well as the encoding, were depicted in a more permissive threshold (*p* < .001 uncorrected) in order to illustrate activations that were obtained on this significance level. The presented coronal slices for the encoding were chosen so that they show anterior (y = −14 mm) and posterior (y = −30 mm) hippocampus (Poppenk, Evensmoen, Moscovitch, & Nadel, 2013).

**Table 1.**
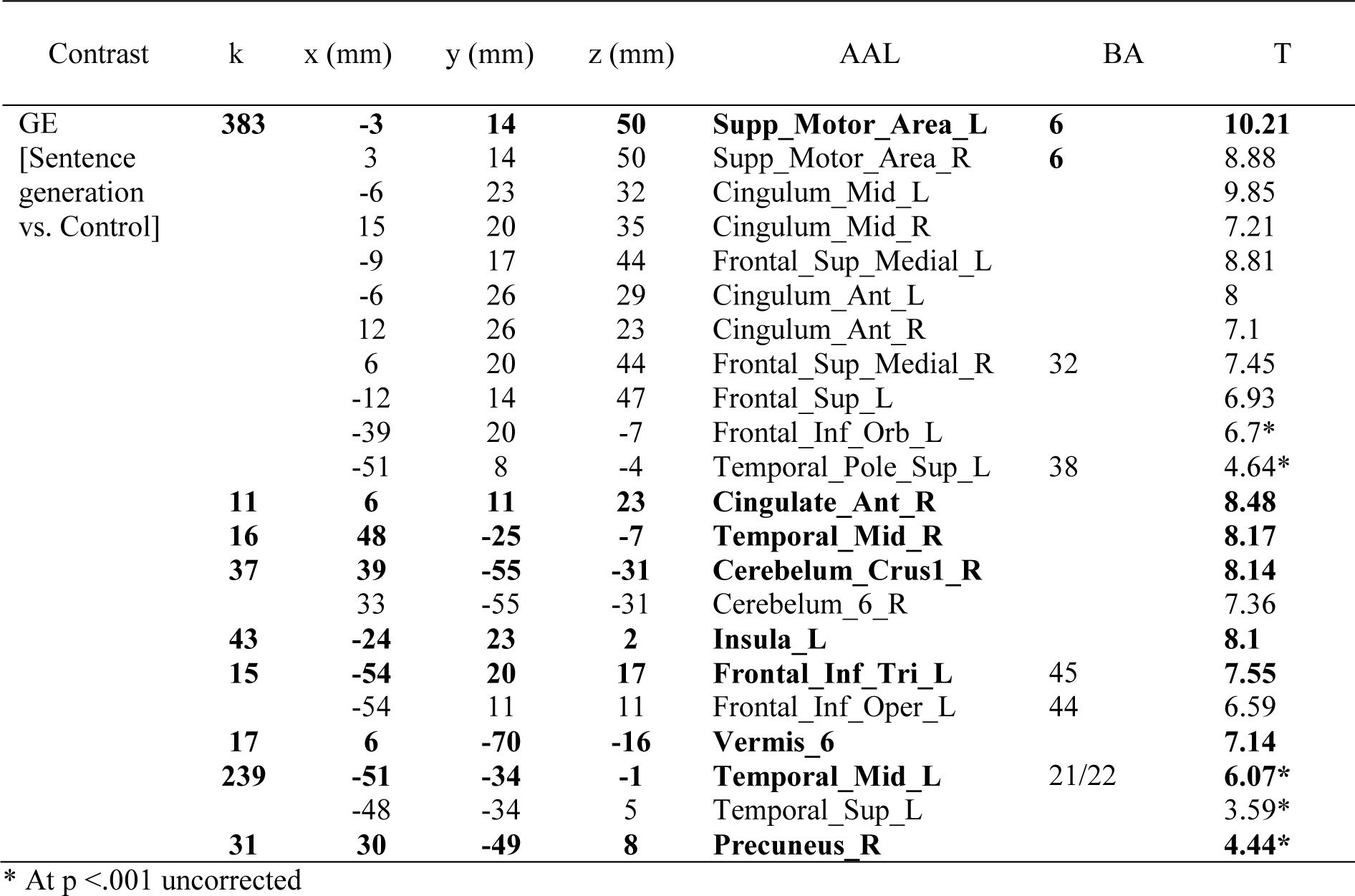
Activated regions for the contrast GE vs. Control. The number of voxels in the cluster (k), the x, y and z coordinates in millimetres, the anatomical region according to AAL atlas (Tzourio-Mazoyer et al., 2002), the Brodmann Area (BA) and the T value are indicated for each peak. All activations were obtained at *p* < .05 (k > 5) corrected except for those with asterisks in the table (\**p* < .001 uncorrected). Abbreviation: GE=sentence Generation with implicit Encoding.

| Contrast | k | x (mm) | y (mm) | z (mm) | AAL | BA | T |
| --- | --- | --- | --- | --- | --- | --- | --- |
| GE | **383** | **-3** | **14** | **50** | **Supp_Motor_Area_L** | **6** | **10.21** |
| [Sentence generation vs. Control] |  | 3 | 14 | 50 | Supp_Motor_Area_R | **6** | 8.88 |
|  |  | -6 | 23 | 32 | Cingulum_Mid_L |  | 9.85 |
|  |  | 15 | 20 | 35 | Cingulum_Mid_R |  | 7.21 |
|  |  | -9 | 17 | 44 | Frontal_Sup_Medial_L |  | 8.81 |
|  |  | -6 | 26 | 29 | Cingulum_Ant_L |  | 8 |
|  |  | 12 | 26 | 23 | Cingulum_Ant_R |  | 7.1 |
|  |  | 6 | 20 | 44 | Frontal_Sup_Medial_R | 32 | 7.45 |
|  |  | -12 | 14 | 47 | Frontal_Sup_L |  | 6.93 |
|  |  | -39 | 20 | -7 | Frontal_Inf_Orb_L |  | 6.7* |
|  |  | -51 | 8 | -4 | Temporal_Pole_Sup_L | 38 | 4.64* |
|  | **11** | **6** | **11** | **23** | **Cingulate_Ant_R** |  | **8.48** |
|  | **16** | **48** | **-25** | **-7** | **Temporal_Mid_R** |  | **8.17** |
|  | **37** | **39** | **-55** | **-31** | **Cerebellum_Crus1_R** |  | **8.14** |
|  |  | 33 | -55 | -31 | Cerebellum_6_R |  | 7.36 |
|  | **43** | **-24** | **23** | **2** | **Insula_L** |  | **8.1** |
|  | **15** | **-54** | **20** | **17** | **Frontal_Inf_Tri_L** | 45 | **7.55** |
|  |  | -54 | 11 | 11 | Frontal_Inf_Oper_L | 44 | 6.59 |
|  | **17** | **6** | **-70** | **-16** | **Vermis_6** |  | **7.14** |
|  | **239** | **-51** | **-34** | **-1** | **Temporal_Mid_L** | 21/22 | **6.07\*** |
|  |  | -48 | -34 | 5 | Temporal_Sup_L |  | 3.59* |
|  | **31** | **30** | **-49** | **8** | **Precuneus_R** |  | **4.44\*** |

\* At $p < .001$ uncorrected

Correct encoding of the items activated the most left anterior hippocampus, bilateral but predominantly left parahippocampus, left prefrontal and middle temporal cortices. Although all of the activation were found on uncorrected threshold (*p* < .001). These activations are presented in the Panel B of the Figure 2 and Table 2.

**Table 2.** Activated regions for the Encoding obtained by parameter modulation of generation activation with respect to correct recognition vs. baseline. The number of voxels in the cluster (k), the x, y and z coordinates in millimetres, the anatomical region according to AAL atlas (Tzourio-Mazoyer et al., 2002), the BA and the T value are indicated for each peak. All activations were obtained at *p* < .001 uncorrected (k > 5).

| Contrast | k | x (mm) | y (mm) | z (mm) | AAL | BA | T |
| --- | --- | --- | --- | --- | --- | --- | --- |
| Encoding<br>[Parameter<br>modulation<br>of generation<br>activation<br>with respect<br>to correct<br>recognition] | **105** | **-30** | **-28** | **-16** | **paraHippocampal L** |  | **5.34\*** |
|  |  | -18 | -13 | -19 | Hippocampus_L | 28 | 5.2* |
|  | **54** | **-36** | **-55** | **17** | **Temporal Mid L** | 22 | **4.77\*** |
|  | 34 | -42 | -73 | 20 | Occipital_Mid_L |  | 4.67* |
|  | **15** | **12** | **-13** | **-22** | **paraHippocampal R** |  | **4.16\*** |
|  | **6** | **-18** | **2** | **53** | **Frontal_Sup_L** |  | **4.15\*** |

\* At $p < .001$ uncorrected

#### 3.2.2. Recognition (RECO)

Retrieval process during the recognition task (task vs. control) activated a large frontal-temporo-parietal network shown in Figure 2, Panel C for old and Panel D for new items and in Table 3 and 4 respectively. The identified network included bilateral fusiform gyri and occipital cortices, left inferior and superior parietal cortices, bilateral cingulum, medial prefrontal cortex, left orbito-frontal gyrus, bilateral insula and bilateral hippocampi and parahippocampi. Both tasks activated bilateral cerebellum VI and recognition of new items activated additionally lobes IV-V and Crus 1.

**Table 3.** Activated regions for the contrast RECO_OLD vs. Control. The number of voxels in the cluster (k), the x, y and z coordinates in millimetres, the anatomical region according to AAL atlas (Tzourio-Mazoyer et al., 2002), the BA and the T value are indicated for each peak. All activations were obtained at *p* < .05 (k > 5) corrected except for those with asterisks in the table (\**p* < .001 uncorrected). Abbreviation: RECO_OLD = recognition of OLD items.

| Contrast | k | x (mm) | y (mm) | z (mm) | AAL | BA | T |
| --- | --- | --- | --- | --- | --- | --- | --- |
| RECO_OLD<br>[OLD vs.<br>Control] | **3030** | **-27** | **-61** | **-16** | **Fusiform_L** |  | **22.45** |
|  |  | 33 | -46 | -22 | Fusiform_R |  | 17.72 |
|  |  | -27 | -49 | -22 | Cerebelum_6_L |  | 19.77 |
|  |  | 30 | -46 | -22 | Cerebelum_6_R |  | 19.69 |
|  |  | -24 | -31 | -13 | ParaHippocampal_L |  | 4.02* |
|  |  | 21 | -28 | -13 | ParaHippocampus_R |  | 4.08* |
|  | **484** | **-39** | **-31** | **44** | **Postcentral_L** | 40 | **16.12** |
|  |  | -30 | -58 | 47 | Parietal_Inf_L |  | 12.04 |
|  |  | -36 | -13 | 59 | Precentral_L |  | 9.09 |
|  |  | -27 | -58 | 50 | Parietal_Sup_L |  | 8.65 |
|  |  | -45 | -31 | 32 | SupraMarginal_L | 40 | 7.61 |
|  | **182** | **-3** | **23** | **44** | **Frontal_Sup_Medial_L** |  | **10.63** |
|  |  | 0 | -4 | 56 | Supp_Motor_Area_L |  | 10.29 |
|  |  | -3 | 11 | 44 | Supp_Motor_Area_L |  | 10.24 |
|  |  | 6 | 17 | 44 | Cingulum_Mid_R |  | 8.86 |
|  |  | -6 | 11 | 41 | Cingulum_Mid_L |  | 7.55 |
|  |  | 3 | 23 | 44 | Frontal_Sup_Medial_R |  | 7.95 |
|  | **67** | **-27** | **20** | **2** | **Insula_L** |  | **9.85** |
|  |  | -33 | 26 | -4 | Frontal_Inf_Orb_L |  | 7.85 |
|  | **32** | **33** | **20** | **2** | **Insula_R** |  | **9.68** |
|  | **30** | **-18** | **-28** | **-1** | **Thalamus_L** |  | **9.62** |
|  |  | -21 | -28 | -7 | Hippocampus_L |  | 6.85 |
|  | **37** | **21** | **-28** | **-4** | **Hippocampus_R** |  | **9.07** |
|  |  | 6 | -28 | -4 | Thalamus_R |  | 7.06 |
|  | **27** | **-57** | **8** | **26** | **Precentral_L** |  | **8.98** |
|  | **15** | **-6** | **-28** | **-7** | **Lingual_L** |  | **8.45** |

\* At $p < .001$ uncorrected

**Table 4.**
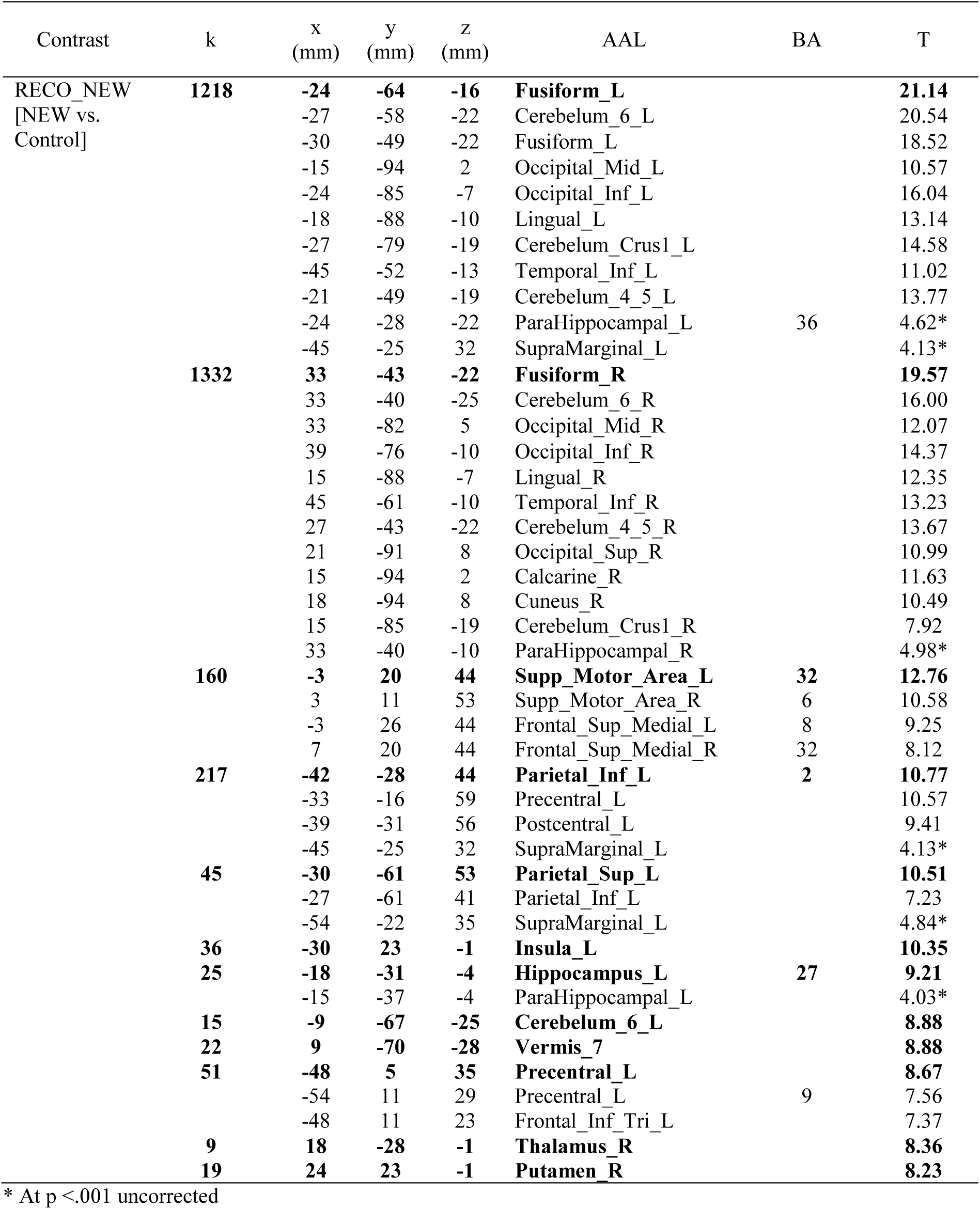
Activated regions for the contrast RECO_NEW vs. Control. The number of voxels in the cluster (k), the x, y and z coordinates in millimetres, the anatomical region according to AAL atlas (Tzourio-Mazoyer et al., 2002), the BA and the T value are indicated for each peak. All activations were obtained at *p* < .05 (k > 5) corrected except for those with asterisks in the table (\**p* < .001 uncorrected). Abbreviation: RECO_NEW = recognition of NEW items.

When compared the recognition of two types of items we could observe that the recognition of old items engaged much more the left parietal cortex, notably precuneus, cuneus and angular gyrus, as well as bilateral middle cingulate and middle temporal cortices. On the other hand, correctly rejecting new items in comparison to correctly recognizing old ones activated more bilateral fusiform and occipital regions. The activations are presented in the Table 5.

**Table 5.** Activated regions for the contrast RECO_OLD vs. RECO_NEW and the opposite contrast. The number of voxels in the cluster (k), the x, y and z coordinates in millimetres, the anatomical region according to AAL atlas (Tzourio-Mazoyer et al., 2002), the BA and the T value are indicated for each peak. All activations were obtained at *p* < .05 (k > 5) corrected except for those with asterisks in the table (\**p* < .001 uncorrected). Abbreviation: RECO_OLD = recognition of OLD items; RECO_NEW = recognition of NEW items.

| Contrast | k | x (mm) | y (mm) | z (mm) | AAL | BA | T |
| --- | --- | --- | --- | --- | --- | --- | --- |
| RECO_OLD vs. RECO_NEW | **210** | **-6** | **-67** | **35** | **Precuneus_L** | **7** | **10.59** |
|  |  | -6 | -64 | 26 | Cuneus_L | 31 | 9.91 |
|  | **31** | **-45** | **-61** | **41** | **Angular_L** |  | **8.47** |
|  | **30** | **3** | **-25** | **35** | **Cingulate_Mid_R** | 24 | **8.10** |
|  |  | -6 | -43 | 35 | Cingulate_Mid_L | 31 | 7.30 |
|  | **8** | **-45** | **-52** | **14** | **Temporal_Mid_L** |  | **7.91** |
|  | **5** | **60** | **-49** | **8** | **Temporal_Mid_R** |  | **7.89** |
| RECO_NEW vs. RECO_OLD | **168** | **27** | **-67** | **-7** | **Fusiform_R** |  | **6.47\*** |
|  |  | 27 | -52 | -10 | Lingual_R |  | 4.33* |
|  | **143** | **-27** | **-85** | **-10** | **Occipital_Inf_L** |  | **6.19\*** |
|  |  | -24 | -88 | -1 | Occipital_Mid_L |  | 5.48* |
|  |  | -27 | -70 | -7 | Fusiform_L |  | 4.87* |
|  | **21** | **-27** | **-49** | **-16** | **Occipital_Mid_L** |  | **4.77\*** |
|  | **11** | **-33** | **-82** | **20** | **Occipital_Mid_R** |  | **4.69\*** |

\* At $p < .001$ uncorrected

#### 3.2.3. Recall (RA)

The recall process (recall vs. baseline) activated a network presented in Figure 2, Panel E and Table 6 that consisted of left inferior frontal and bilateral predominantly right oriented prefrontal and medial frontal cortices and left insula. In addition, bilateral activations in temporal superior and middle cortices as well as left temporal pole were identified. The activation of the parietal regions consisted of the left inferior parietal and angular gyrus, while the activations of the cerebellum were limited to right Crus 1. Right hippocampal activation was also detected when employing lower significance level (*p* < .001).

**Table 6.**
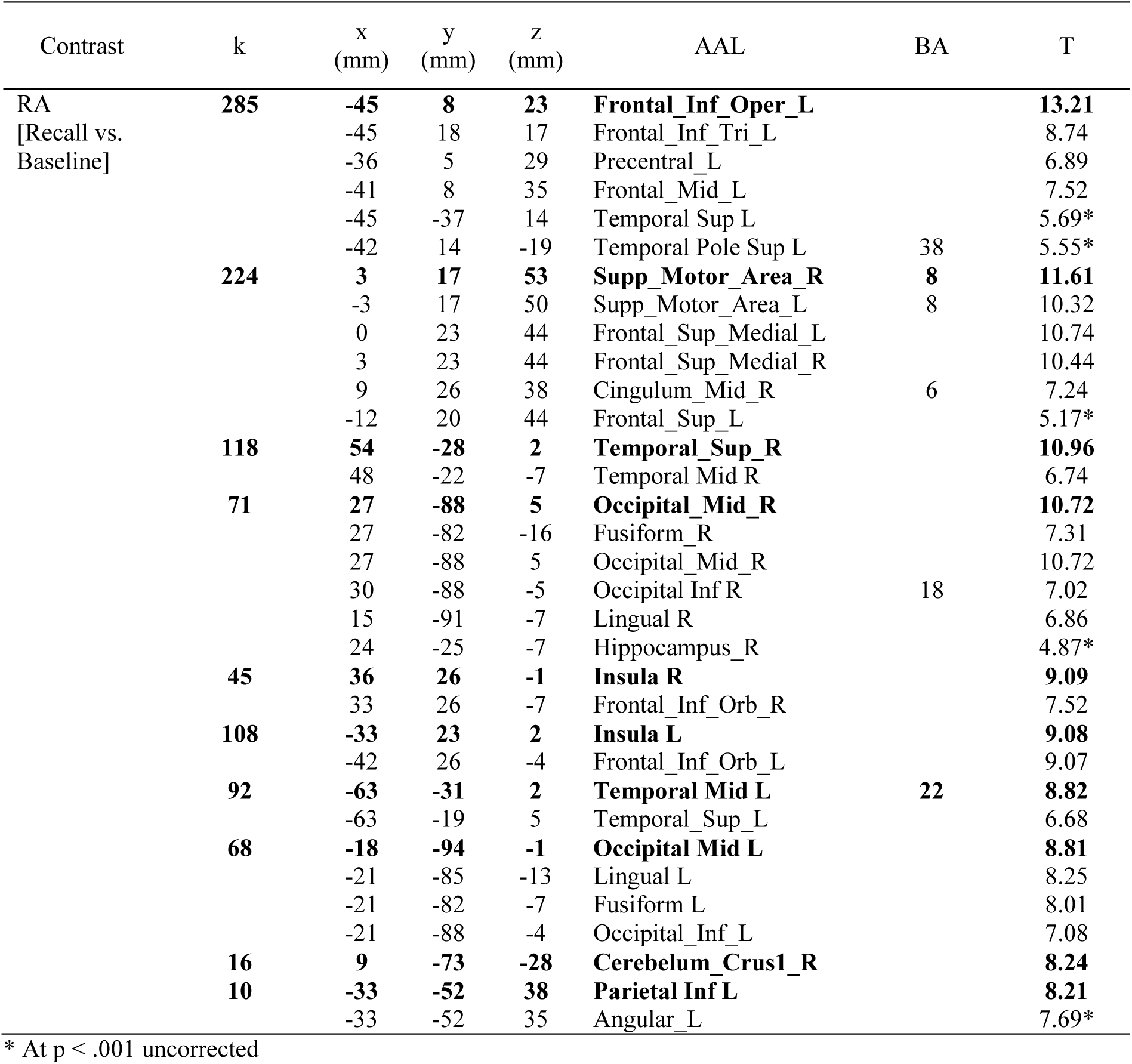
Activated regions for the contrast RA vs. baseline. The number of voxels in the cluster (k), the x, y and z coordinates in millimetres, the anatomical region according to AAL atlas (Tzourio-Mazoyer et al., 2002), the BA and the T value are indicated for each peak. All activations were obtained at *p* < .05 (k > 5) corrected except for those with asterisks in the table (\**p* < .001 uncorrected). Abbreviation: RA = recall.

| Contrast | k | x<br>(mm) | y<br>(mm) | z<br>(mm) | AAL | BA | T |
| --- | --- | --- | --- | --- | --- | --- | --- |
| RA<br>[Recall vs.<br>Baseline] | **285** | **-45** | **8** | **23** | **Frontal_Inf_Oper_L** |  | **13.21** |
|  |  | -45 | 18 | 17 | Frontal_Inf_Tri_L |  | 8.74 |
|  |  | -36 | 5 | 29 | Precentral_L |  | 6.89 |
|  |  | -41 | 8 | 35 | Frontal_Mid_L |  | 7.52 |
|  |  | -45 | -37 | 14 | Temporal Sup L |  | 5.69* |
|  |  | -42 | 14 | -19 | Temporal Pole Sup L | 38 | 5.55* |
|  | **224** | **3** | **17** | **53** | **Supp_Motor_Area_R** | **8** | **11.61** |
|  |  | -3 | 17 | 50 | Supp_Motor_Area_L | 8 | 10.32 |
|  |  | 0 | 23 | 44 | Frontal_Sup_Medial_L |  | 10.74 |
|  |  | 3 | 23 | 44 | Frontal_Sup_Medial_R |  | 10.44 |
|  |  | 9 | 26 | 38 | Cingulum_Mid_R | 6 | 7.24 |
|  |  | -12 | 20 | 44 | Frontal_Sup_L |  | 5.17* |
|  | **118** | **54** | **-28** | **2** | **Temporal_Sup_R** |  | **10.96** |
|  |  | 48 | -22 | -7 | Temporal Mid R |  | 6.74 |
|  | **71** | **27** | **-88** | **5** | **Occipital_Mid_R** |  | **10.72** |
|  |  | 27 | -82 | -16 | Fusiform_R |  | 7.31 |
|  |  | 27 | -88 | 5 | Occipital_Mid_R |  | 10.72 |
|  |  | 30 | -88 | -5 | Occipital Inf R | 18 | 7.02 |
|  |  | 15 | -91 | -7 | Lingual R |  | 6.86 |
|  |  | 24 | -25 | -7 | Hippocampus_R |  | 4.87* |
|  | **45** | **36** | **26** | **-1** | **Insula R** |  | **9.09** |
|  |  | 33 | 26 | -7 | Frontal_Inf_Orb_R |  | 7.52 |
|  | **108** | **-33** | **23** | **2** | **Insula L** |  | **9.08** |
|  |  | -42 | 26 | -4 | Frontal_Inf_Orb_L |  | 9.07 |
|  | **92** | **-63** | **-31** | **2** | **Temporal Mid L** | **22** | **8.82** |
|  |  | -63 | -19 | 5 | Temporal_Sup_L |  | 6.68 |
|  | **68** | **-18** | **-94** | **-1** | **Occipital Mid L** |  | **8.81** |
|  |  | -21 | -85 | -13 | Lingual L |  | 8.25 |
|  |  | -21 | -82 | -7 | Fusiform L |  | 8.01 |
|  |  | -21 | -88 | -4 | Occipital_Inf_L |  | 7.08 |
|  | **16** | **9** | **-73** | **-28** | **Cerebelum_Crus1_R** |  | **8.24** |
|  | **10** | **-33** | **-52** | **38** | **Parietal Inf L** |  | **8.21** |
|  |  | -33 | -52 | 35 | Angular_L |  | 7.69* |

\* At $p < .001$ uncorrected

Figure 3 presents the synthesis of the results. The principal findings can be summed up as follows: (a) sentence generation activated bilateral temporal and left frontal and parietal regions, (b) implicit encoding of the items into the long-term memory engaged bilateral (para)hippocampal and left prefrontal activations, (c) recognition of the items activated bilateral inferior ocipito-temporal, left parietal and bilateral hippocampal and parahippocampal regions, but also the left frontal inferior, bilateral SMA and (d) recall that assessed the language-and-memory interaction activated large fronto-temporo-parietal network with right hippocampus.

**Figure 3.**
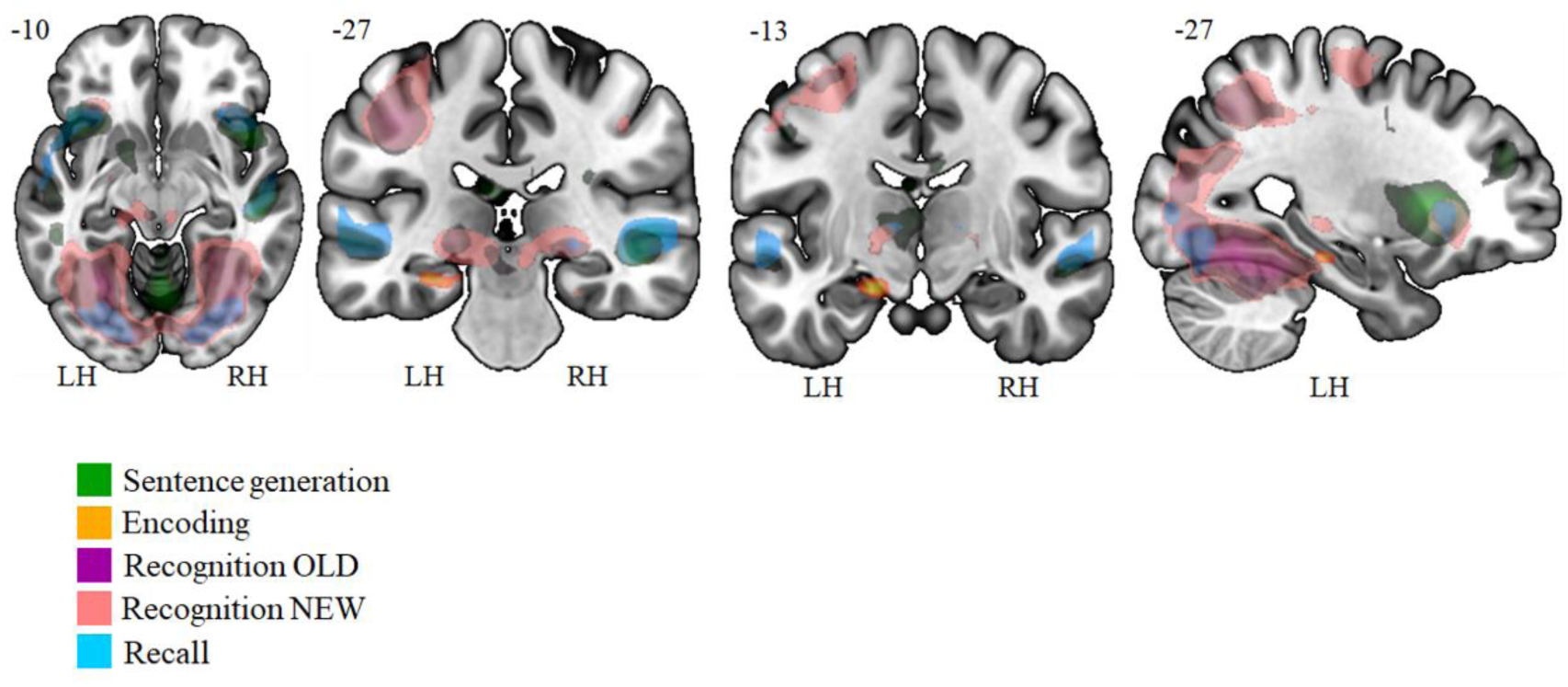
Illustrative overview of the synthesis of results obtained with GE2REC protocol during sentence generation (green), encoding (orange), recognition of old (violet) and new items (pink) as well as the recall (blue). The activated regions are projected onto 2D anatomical slices presented in axial, coronal and sagittal orientations. The left (LH) and right (RH) hemispheres are indicated.

## 4. Discussion

The interaction between language and memory plays an essential role in our everyday lives, the most obvious example being that it allows us to hold conversations; memory provides the basis of understanding we need to track and maintain proper conversational flow. This interaction is equally observable, for example, when lesions occur in language regions, but symptoms appear in the memory domain or vice versa, a frequent scenario in several neurological, neuropsychological and psychiatric disorders, such as post-stroke aphasia (Schuchard and Thompson, 2014), temporal lobe epilepsy (Jaimes-Bautista et al., 2015; Everts et al., 2010), conditions that cause auditory hallucinations (Ćurčić-Blake et al., 2017), and even certain language problems were found in the case of H.M. (MacKay, Stewart & Burke, 1998). We know less about the specific cerebral substrate underlying language and memory interaction and how the two functions are able to dynamically modulate each other. These questions are fundamental from a neurocognitive standpoint given that cognitive functions are no longer considered as monolithic with modular representation, but instead are linked by their interaction based on key common brain substrates. In order to explore these matters, we need an adequate tool that would be able to capture this synergy in action while being adapted to both clinical setting and standards in empirical research. As a reaction to that need, we have developed the GE2REC protocol that interactively maps the language-and-memory network. The present study provides its evaluation in healthy participants. In the following we will first discuss each run of GE2REC separately, demonstrating at the end the joint language-and-memory network cartography it provides.

Our results indicate that sentence generation activated a large bilateral, but predominantly left fronto-temporo-parietal network (Figure 2 and Table 1). This network is in line with the typical representation of language, as suggested by current models (Hickok & Poeppel, 2007; Indefrey & Levelt, 2004; Price, 2012) and a meta-analysis (Vigneau et al., 2006). The network includes left inferior frontal (pars opercularis and pars triangularis), left insula and bilateral SMA usually required by the production of sentences (Grande et al., 2012; Haller, Radue, Erb, Grodd, & Kircher, 2005; Menenti, Segaert, & Hagoort, 2012; Price, 2012; Segaert, Menenti, Weber, Petersson, & Hagoort, 2012). This activation can be explained by a significant planning, articulatory and syntactic processes required by GE2REC, even if the language was produced covertly (Benjamin et al., 2017; Haag & Bonelli, 2013). Regarding the temporo-parietal network, superior and middle temporal gyri as well as the superior temporal pole were activated, which is in line with other results reporting syntactic (Grande et al., 2012; Menenti et al., 2012; Segaert et al., 2012), lexico-semantic (Grande et al., 2012; Menenti et al., 2012; Price et al., 2012) and phonological (Indefrey & Levelt, 2004; Price, 2012) demands during a sentence generation. The covert form of the generation might indeed involve inhibitory processes (of overt language with articulation and vocalization) explaining the activation of anterior cingulum (Loevenbruck et al., 2018; Price, 2012). Nevertheless, the sentence generation was not purely language task but it also included implicit encoding. The obtained activations (Figure 2 and Table 2), although all at a more permissive threshold (*p* < .001), were in line with previously reported findings for successful encoding such as bilateral hippocampi and right parahippocampus, together with left frontal superior, as reported by models (Diana et al., 2007; Preston & Eichenbaum, 2013) and a meta-analysis (Spaniol et al., 2009). The activations were predominantly left lateralized, explained by the encoding processing itself (Spaniol et al., 2009) and the use of a verbal material (Gabrieli, 2001), even if according to HERA model, the left prefrontal cortex should be involved, regardless the material type (Habib, Nyberg, & Tulving, 2003). Along an anterior-posterior axis of the hippocampus, relying on the anterior-posterior distinction coordinates proposed by Poppenk and his colleagues (2013), the obtained encoding activation tended to be rostral (anterior), in line with previous studies and models (Lepage et al, 1998; Preston & Eichenbaum, 2013; Spaniol et al., 2009).

The change in the modality between GE (auditory) and RECO (visual) run was implemented in this protocol with the intention of eliciting participants’ responses based on recognition, rather than familiarity, activating thus episodic memory (Perrone-Bertolotti et al., 2015). We believe our participants really did remember instead of relied on familiarity of the stimuli since the network activated by RECO corresponded well with the “Binding of Item and Context” model (BIC, Diana et al., 2007) and the episodic posterior medial network proposed by Ranganath and Ritchey (2012) in that it indeed activated posterior bilateral hippocampal and parahippocampal gyri, as well as cingulate, lateral parietal and medial prefrontal cortices (as a part of the default mode network, DMN). We obtained bilateral instead of right prefrontal activity predicted by the HERA model (Habib et al., 2003), our result being in agreement with other studies (Spaniol et al., 2009). Additionally, the differences we found between correctly identified old items and correctly rejected new items reflected in reaction time and left parietal activation are in line with previous studies (Guerin & Miller, 2009). Although previous studies found engagement of fusiform gyrus, inferior frontal cortex and insula during both encoding and retrieval processes (Aldenkamp et al., 2003; Spaniol et al., 2009), we believe that the activations found during recognition may reflect a verbal strategy used by participants to perform the task and that they engaged in picture naming. Namely, inferior frontal, SMA, insula, fusiform and parietal cortices correspond well with the picture naming network (Duffau et al., 2014). Additionally, cerebellar activations we found, specifically Crus 1 and lobules IV-V and VI, correspond to language processes demanded by the task (Keren-Happuch et al., 2014; Price, 2012; Stoodley & Schmahmann, 2018). These results underline, in addition to the specificity of our task, a real difficulty to disentangle language and memory processes. Trying to separate these two functions is probably artificial, instead cognitive functions should be assessed realistically in a dynamic interaction (Kellermann et al., 2016), especially when it comes to patients. Indeed, having in mind for instance that TLE is often accompanied by HS with implications on both language and memory (Alessio et al., 2006; Bonelli et al., 2011; Davies et al., 1998; Zalonis et al., 2017), it is crucial that the protocol used in preoperative mapping has the ability to robustly activate hippocampal and neighbouring structures. We have seen that the GE2REC protocol can activate these structures both during encoding and recognition memory processes. Still, the validation of activation of mesial temporal lobe structures should be further tested by detailed delineation of these regions (de Vanssay-Maigne et al., 2011) which goes beyond the scope of the current study.

Finally, the RA task was designed so that it assesses directly the interactive dynamics of the language-and-memory but also to resemble everyday life experience, by having a more natural recollection context. The ecological dimension of tasks used in patients is, indeed, particularly important (Mayer & Murray, 2003). Bearing this goal in mind, one of the important features of the GE2REC protocol is its proximity to everyday experiences, which is most evident in the RA task. The identified network during this task resembled the one found in the generation task that included the activations of the left inferior frontal gyrus, bilateral SMA and insula as well as bilateral superior and middle temporal cortices and Crus 1 of the cerebellum. These activations can be related to the language component of the network (Hickok & Poeppel, 2007; Indefrey & Levelt, 2004; Price, 2012). On the other hand, the activations of the bilateral prefrontal and predominantly left parietal cortices as well as bilateral fusiform gyri, are in agreement with the previous results on memory retrieval (Aldenkamp et al., 2003; Spaniol et al., 2009). It should also be noted that some structures that were active during this task have previously been found to be active both in language and memory tasks. For example, temporo-polar cortex, lateral orbitofrontal and angular gyrus make up a part of the two memory systems proposed by Ranganath and Ritchey (2012), while at the same time being involved in language networks and engaged in semantic processing (Duffau et al., 2014; Price, 2012). Additionally, the occipito-temporal, parietal and hippocampal activations we found during the RA task match the subsystem identified by Vandenberghe and his colleagues (2013) who interpreted it as a link between inner representations and episodic memory. This again supports the idea of a large language-and-memory network and shows that these regions are activated when the individual is engaged in mixed language-and-memory tasks and situations. Importantly, although hippocampus was proposed to be included in language network (Covington & Duff, 2016) being found to have role in different language tasks, notably retrieving lexically and semantically associated words, naming, language comprehension tasks and syntactic processing (Aldenkamp et al., 2003; Meyer et al., 2005; Tracy & Boswell, 2008), we observed its activation only during the RECO and RA task but not the GE. Nevertheless, this is not evidence that refutes the implication of the hippocampus in language processes; on the contrary, there are several explanations for the lack of activations. First of all, it could be that hippocampus is implied in other aspects of language processing that we have not included in the GE task such as sentence processing (Piai et al., 2016), while it was active during picture naming that we assume was performed during RECO task. Secondly, it has been proposed that comprehension of familiar words (such as those used in our protocol) activate nodes that have already formed connections, so there is no need for new connection formation and hippocampal activity (MacKay et al., 1998). Finally, our results could also suggest that hippocampus is perhaps not a primary element of the exclusive language network, but that it is instead a part of the language-and-memory network, connecting the two systems. Overall, the wide network (Figure 3) recruited by the GE2REC protocol, can be considered as the interactive language-and-memory network since it was obtained through the linked tasks in which two processes were highly intertwined. It is also important to note that it is not merely a provisional sum of activations, but it is the cerebral substrate of combined and intermixed language and memory processes and has specific anatomical support. Specifically, the mesial temporal, temporal pole and prefrontal cortices could be inter-connected via the direct inter-hippocampal pathway, while the polysynaptic pathway could connect parietal and temporal cortices through the parahippocampal gyrus towards cingulate cortices, as suggested by Duvernoy (2005). Additionally, anterior temporal and orbito-frontal areas that have been found during RA could be connected via UF that supports both functions (Diehl et al., 2008; Duffau, 2014; Kucukboyaci et al.,2014; McDonald et al., 2008). IFOF could connect frontal and occipital regions, supporting semantic processing, verbal memory and noetic consciousness (McDonald et al., 2008; Moritz-Gasser et al., 2013). Nevertheless, one of the next step of this work is to explore structural and functional connectivity within the GE2REC language-and-memory network.

Our work has several limitations which we will discuss in the next lines. First of all, participants’ responses for GE and RA task cannot be recorded since the covert speech was used. Consequently, we are not able to measure performance of these tasks. Nevertheless, as well as previous studies employing the covert instead of overt response modality (Benjamin et al., 2017; Haag & Bonelli, 2013), we too identified expected cognitive networks. Secondly, participants’ responses during RECO were not so highly accurate (but clearly far above the chance level) as expected and as shown previously (Perrone-Bertolotti et al., 2015). This can be explained by the specificity of GE2REC tasks. Indeed, participants were not explicitly instructed to memorize the item they heard during GE. In addition, drawing on the important capacity of episodic memory to flexibly retrieve and recombine information from distinct past experiences (Carpenter & Schacter, 2017) and given that the images used during RECO were rather familiar (e.g. bottle, orange etc.) and frequently encountered in the everyday life, participants could have mistakenly combined features of different episodes. Finally, it should be noted that this protocol does not aspire towards general and exhaustive models of language and memory because many linguistic and memory aspects are not explored by GE2REC.

## 5. Conclusion

We proposed the GE2REC protocol that we tested in the current study in healthy participants, to interactively map a global language-and-memory network. The main goal was to propose a protocol that could provide significant information on the neurocognition of language and memory. GE2REC is also easy to perform, has short duration and sufficient robustness of the activation. GE2REC protocol can jointly activate a large fronto-temporo-parietal network generally observed in language studies, as well as mesial temporal, parietal and prefrontal cortices, generally reported by memory studies. In addition, with respect to the memory, it explores both encoding and retrieval processes and allows for left-right and anterior-posterior segregation of their cerebral representations. By synthesizing the results of its three tasks designed to explore the interactive nature of language-and-memory, GE2REC provides the cartography of this network which could be of practical importance.

## Acknowledgments

IRMaGe MRI/Neurophysiology facility was partly funded by the French program “Investissement d’Avenir” run by the “Agence Nationale pour la Recherche”; grant “Infrastructure d’avenir en Biologie Santé” [grant number ANR-11-INBS-0006].

## Footnotes

1 We decided to use different, real images rather than original ones from DO80 test, for two reasons. First, the original images are black-and-white drawings and taking the real images of objects was closer to everyday experience. And more importantly, DO80 is frequently used in behavioral experiments and neuropsychological evaluation of patients in our environment. Thus, we wanted to avoid the recognition of items based on familiarity.

